# A precise balance of TETRASPANIN1/TORNADO2 activity is required for vascular proliferation and ground tissue patterning in Arabidopsis

**DOI:** 10.1101/2023.10.18.562921

**Authors:** Nataliia Konstantinova, Eliana Mor, Eline Verhelst, Jonah Nolf, Kenzo Vereecken, Feng Wang, Daniel Van Damme, Bert De Rybel, Matouš Glanc

## Abstract

The molecular mechanisms guiding oriented cell divisions in the root vascular tissues of *Arabidopsis thaliana* are still poorly characterized. By overlapping bulk and single-cell transcriptomic datasets, we unveiled *TETRASPANIN1 (TET1)* as a putative regulator in this process. *TET1* is expressed in root vascular cells and loss-of-function mutants contain fewer vascular cells files. We further generated and characterized a CRISPR deletion mutant and show, unlike previously described mutants, that the full knock out is additionally missing endodermal cells in a stochastic way. Finally, we show that HA-tagged versions of TET1 are functional in contrast to fluorescent TET1 translational fusions. Immunostaining using HA-TET1 lines complementing the mutant phenotype revealed a dual plasma membrane and intracellular localization in the root vasculature and a polar membrane localization in young cortex, endodermal and initial cells. Taken together, we show that TET1 is involved in both vascular proliferation and ground tissue patterning. Our initial results pave the way for future work into deciphering its precise mode of action.

**Summary statement:** This study reveals a novel role of tetraspanin TET1/TRN2 in root vascular development and ground tissue patterning in the model plant *Arabidopsis thaliana*.

## Introduction

Oriented cell divisions are integral to the process of creating the three-dimensional shape of a plant (Metcalfe and Esau, 1966). This is particularly evident in the primary root apical meristem (RAM) of *Arabidopsis thaliana* (Arabidopsis), where cells are organized in a highly conserved pattern of concentric layers, achieved via strict spatiotemporal control of cell division orientation (Dolan et al., 1993). Anticlinal (perpendicular to the root surface and growth axis) division plane positioning adds cells to existing cell files and leads to longitudinal growth; whereas radial (perpendicular to the surface but parallel to the axis) and periclinal (parallel to both the surface and the growth axis) divisions increase the number of cell files (Smet and De Rybel, 2016). Under normal growth conditions, epidermal and ground tissues undergo periclinal divisions in the stem cell niche: the protoderm initials give rise to lateral root cap (LRC) and epidermis cell files, while cortex-endodermis initial (CEIs) daughter cells (CEIDs) produce cortex and endodermis. Upon exiting the stem cell region, LRC, epidermis, cortex and endodermis cells undergo anticlinal divisions while they retain the capacity to switch the division plane upon e.g. wounding (Marhavá et al., 2019) or ectopic expression of specific transcription factors (TFs) (De Rybel et al., 2013; Miyashima et al., 2019; Ohashi-Ito et al., 2014; Smet et al., 2019). The situation is different for vascular cells, as radial and periclinal divisions are needed during e.g. the formation of the phloem lineage during primary growth (Miyashima et al., 2019; Rodriguez-Villalon et al., 2014).

Cell division orientation in the vasculature is controlled by a complex transcriptional network. One key regulator is the basic helix-loop-helix (bHLH) TF heterodimer formed by TARGET OF MONOPTEROS5 (TMO5) and LONESOME HIGHWAY (LHW). Upon misexpression, this complex is capable of triggering ectopic periclinal/radial divisions in any cell type (De Rybel et al., 2013; De Rybel et al., 2014; Katayama et al., 2015; Mor et al., 2022; Ohashi-Ito et al., 2014). Several factors acting both upstream and downstream of TMO5/LHW have been described (Vera-Sirera et al., 2015; Wybouw et al., 2023; Yang et al., 2021). Notably the DNA-BINDING WITH ONE ZINC FINGER2.1 (DOF2.1) TF, a direct TMO5/LHW target that is sufficient to trigger ectopic periclinal radial divisions and is related to the PEAR (PHLOEM EARLY DOF) TFs that control vascular cell division orientation independently of TMO5/LHW (Miyashima et al., 2019; Smet et al., 2019). Nonetheless, the post-transcriptional mechanisms involved in executing the switch in cell division plane positioning downstream of these transcriptional programs remain poorly understood.

Division plane formation is often viewed as a transient polarization event and is as such expected to depend on similar mechanisms as other polarity establishment pathways (Glanc, 2022; Müller, 2019). Cell polarization in plant and non-plant systems typically involves positive feedback loops that mechanistically rely on protein scaffolding (Chiou et al., 2017; Ramalho et al., 2021). Indeed, polarly localized scaffolding proteins, such as BASL, POLAR and SOSEKI proteins, have been shown to drive cell division processes in a polarized manner (Dong et al., 2009; Houbaert et al., 2018; Yoshida et al., 2019). In this work, we investigate the involvement of TETRASPANIN1/ TORNADO2 (TET1/TRN2), a transmembrane protein with a putative scaffolding function, in the regulation of cell division orientation during vascular development.

TET1 belongs to the TETRASPANIN (TET) protein family, which is conserved across multicellular eukaryotes and is represented by 17 members in Arabidopsis (Boavida et al., 2013; Wang et al., 2015; Wang et al., 2012). TETs are small membrane proteins with a conserved fold consisting of four membrane-spanning helices connected by three loops: a small extracellular, a small intracellular and a large extracellular loop (Reimann et al., 2017). TETs function in a variety of cellular and developmental processes in animal systems, ranging from endomembrane trafficking to signalling and cell adhesion (Charrin et al., 2014; Reimann et al., 2017). Scaffolding and partitioning of various multiprotein complexes into the so-called TETRASPANIN-Enriched Microdomains (TEMs) at the plasma membrane (PM) is a recurrent mode of TET action across their diverse functional roles (Reimann et al., 2017). Comparatively, less is characterized when it comes to the function and localization of TETs in plants, although it has been shown that their PM localization depends on the presence of a “GCCK/RP” motifs and cysteine residues (Zhu et al., 2022). Most single mutants show no striking phenotypes, suggesting a high degree of functional redundancy in the family (Wang et al., 2015). One exception is the *tet1/trn2* single mutant, which shows prominent pleiotropic defects ranging from dwarfed overall architecture, to sterility and asymmetrical venation patterning (Cnops et al., 2000; Cnops et al.,2006; Olmos et al., 2003; Chiu et al., 2007). Additionally, the *tet13* mutant shows a shortened meristem and reduced emergence of lateral roots, suggesting that TET13 plays a role in formation of the primary root (Wang et al., 2015). TET8 and TET9 contribute to plant immunity, as double mutants are susceptible to *Botrytis cinerea*. These double mutants also have a reduction of host sRNA in the excreted exosomes, required for silencing of fungal genes essential for infection, while the single *tet8* mutant secretes less extracellular vesicles (Cai et al., 2018; He et al., 2021; Liu et al., 2020). TET5 and TET6 are reported to have a role in plant growth, as the *tet5 tet6* double mutant has larger leaves and a longer primary root (Wang et al., 2015). Although it is thus clear that TETs are involved in a wide range of developmental aspects, a thorough understanding of their functionality is still lacking. In this study, we characterize TET1 in the context of vascular development, using cell biological and genetic approaches. We demonstrate that both the *tet1* mutant phenotypes and the *TET1* expression pattern argue for a role in the maintenance of root vascular tissues and ground tissue patterning. In addition, we show that an optimal balance of *TET1* is required for proper growth and suggest that the mode-of-action might be linked to intracellular trafficking events.

## Results

### Identification of TET1 as regulator of division orientation in vascular cells

To identify novel regulators of cell division orientation during vascular development, we harnessed a published single-cell RNA sequencing (scRNA-seq)-based transcriptomic atlas of the Arabidopsis root (Wendrich et al., 2020). Specifically, we focused on transcripts enriched in young protophloem and/or procambium cells, which undergo a switch from anticlinal to periclinal or radial cell division orientation most often outside of the stem cell niche (Miyashima et al., 2019). We further reasoned that key factors involved in vascular division plane positioning would likely act downstream of the TMO5/LHW-DOF2.1 pathway (De Rybel et al., 2013; Smet et al., 2019). With this rationale, we transcriptionally profiled the p*RPS5A::DOF2.1-GR* inducible misexpression line (Smet et al., 2019) in a time-course induction using bulk RNA-seq (**Table S1**) and overlapped the transcripts upregulated upon *DOF2.1* induction with the phloem/procambium-enriched candidates obtained from the scRNA-seq dataset (Wendrich et al., 2020) (**Table S1**, **Figure 1A**). To further narrow down the search space, we focused on genes linked to entities involved in the execution of cell division, such as the cytoskeleton, plasma membrane or the cell wall (Glanc, 2022; Livanos and Müller, 2019). Among the shortlist of candidate genes identified by this approach, *TETRASPANIN1/ TORNADO2 (TET1/TRN2)* drew our attention for several reasons. Firstly, besides its enrichment in young protophloem, procambium and the stem cell niche, lower amounts of the *TET1* transcript were detected in most other cell types as well, which would be expected for a regulator of core cellular process such as cell division (**Figure S1**). Secondly, *TET1* was consistently and significantly upregulated upon DOF2.1-GR induction from 1h after DEX treatment onwards (**Table S1**) and *tet1/trn2* mutants have strong pleiotropic growth and patterning defects which include lateral root cap (LRC)/epidermis cell type mis-specification, presumably caused by irregular protoderm initial cell divisions (Cnops et al., 2000; Cnops et al., 2006; Lieber et al., 2011; Olmos et al., 2003; Yamaguchi et al., 2017). Thirdly and most important, TET proteins are conserved across multicellular eukaryotes and function as molecular scaffolds in a plethora of multiprotein complexes at the plasma membrane in animal cells (Reimann et al., 2017; Termini and Gillette, 2017). As protein scaffolding is a common feature of numerous cellular polarity establishment pathways (Ramalho et al., 2021), this provides a mechanistically plausible role for TET1 in plant cell division orientation, which is thought to be tightly linked to cell polarity (Glanc, 2022; Müller, 2019). In summary, based on overlap analysis of two different transcriptomic datasets, we selected TET1 as a putative regulator of cell division orientation during vascular development.

**Figure 1:**
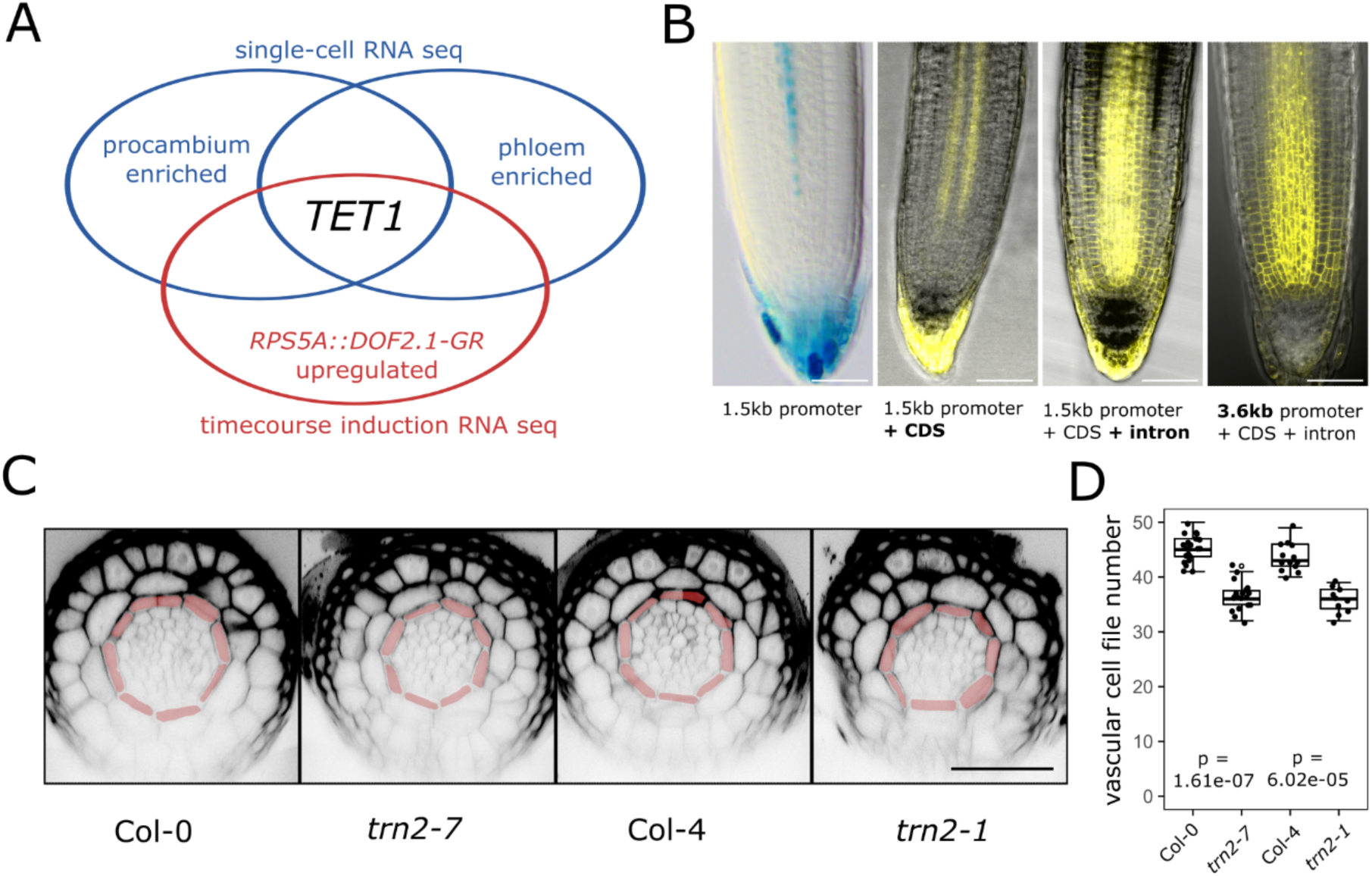
TET1 expression pattern and vascular *tet1* mutant phenotype. **A)** Schematic of the overlap analysis of the inducible *pRPS5A::DOF2.1* RNA-seq and the root single cell atlas dataset that lead to identification of *TET1*. **B)** Left to right: longitudinal section of a GUS stained *pTET1::nlsGUS-GFP,* confocal image of *pTET1_1500_::TET1-sYFP, pTET1_1500_::TET1g-sYFP,* and *pTET1_3600_::TET1g-sYFP,* with bright field overlay. Scale bar = 50µM. **C)** Representative images of transverse cross-sections of PS-PI stained meristematic roots of Col-0, *trn2-7,* Col-4, and *trn2-1,* used for quantification shown in panel d. Red false colouring represents endodermis cell layer. Scale bar = 50µM. **D)** Quantification of the vascular cell file number for the plant lines Col-0, *trn2-7,* Col-4, and *trn2-1*.

Although the scRNA-seq data (Wendrich et al., 2020) suggested mRNA expression of *TET1* in most cell types of the root meristem with an enrichment in young protophloem and procambium (**Figure S1**), this is contradicted by the expression pattern of the previously described *pTET1::nlsGUS-GFP* transcriptional reporter line (Wang et al., 2015). This line reporting on *TET1* promoter activity revealed weak expression in a small subset of vascular and root cap cells and only in lines with multiple T-DNA insertions (Wang et al., 2015). We thus hypothesized that endogenous *TET1* expression depended on additional regulatory elements in a longer promoter region than used previously and/or the protein coding region. To test this hypothesis, we generated a series of native promoter-driven translational reporter constructs varying in the length of the promoter fragment and presence/absence of the single 595 bp intron. The expression pattern of the cDNA-derived *pTET1_1500_::TET1-sYFP* was very similar to the previously described *pTET1::nlsGUS-GFP*, while the gDNA-derived, intron-including *pTET1_1500_::TET1g-sYFP* showed a comparably stronger and broader expression, corresponding to the predicted scRNA-seq data (**Figure 1B**). To test whether *TET1* expression in the root meristem is regulated by additional elements further upstream in the promoter region, similarly to the situation in developing flowers (Yamaguchi et al., 2017), we generated a *pTET1_3600_::TET1g-mCitrine* construct using a 3600 bp promoter fragment spanning the whole region between the start codon of *TET1* and the stop codon of the 5’ neighbouring gene, *bHLH071*. In the root meristem, the expression patterns of *pTET1_1500_::TETg1-sYFP* and *pTET1_3600_::TET1g-mCitrine* were indistinguishable (**Figure 1B**), indicating that while the longer promoter fragment is important for correct *TET1* expression in flowers and perhaps other tissues, expression in the root meristem depends primarily on regulatory elements present in the intron.

To further validate the role of TET1 in the regulation of vascular development, we probed the known *tet1/trn2* mutants for defects in the anatomy of the root apical meristem vascular bundle. Specifically, we analysed the number of vascular cell files, which is a reliable readout of the relative frequency of periclinal/radial cell divisions (Arents et al., 2022; De Rybel et al., 2013; Smet et al., 2019). At least 7 different *tet1* alleles including T-DNA insertion lines and mutants obtained via EMS mutagenesis have been described to date; all causing the characteristic *tornado* phenotype which includes dwarfism, sterility and severe twisting of roots, leaves and other organs (Cnops et al., 2000; Cnops et al., 2006; Lieber et al., 2011; Olmos et al., 2003; Yamaguchi et al., 2017). The root vascular phenotype of *tet1/trn2* mutants has however not been examined so far. We thus quantified the number of cell files within the vascular bundle in the primary root meristem of two different *tet1* alleles: *trn2-1* (in the Col-4 background) and *trn2-7* (in the Col-0 background). In both cases, we observed a significant reduction in vascular cell file number compared to the respective wildtype control (**Figure 1C,D** and **Supplemental Table S5**). This result indicated a lower frequency of a division plane switch from anticlinal to periclinal/radial during the development of primary root vasculature in *tet1* mutants. Collectively, the *TET1* expression pattern and loss-of-function mutant phenotype strongly support a role for TET1 during root vascular proliferation, presumably in the control of cell division orientation.

### Generation of a *tet1* deletion mutant reveals both vascular and ground tissue defects

To get an insight into a possible molecular mechanism for TET1 function, we next aimed to determine its subcellular localization. To avoid artifacts caused by non-functional reporters, we first tested the functionality of our fluorescent protein fusions (**Figure 1B**) by a genetic complementation assay. To this end, we transformed the constructs into *trn2-7* heterozygote plants, as all known homozygous *tet1* mutants are sterile (Cnops et al., 2000; Cnops et al., 2006). After screening T1 and T2 populations of 13 different TET1 translational reporter constructs, consisting of different combinations of endogenous and constitutive promoters, gDNA and cDNA fragments and different N- and C-terminal tags (**Table S2**), we consistently observed ∼25% plants with the characteristic *tornado* phenotype (**Figure S2**), and never found a *tet1/trn2-7* homozygote with a wildtype-like phenotype. This indicated that none of the 13 tested constructs rescued the *tet1/trn2* mutant. Such lack of rescue could in theory be caused by an incorrect expression pattern of the transgenes or non-functional fusion proteins. Second-site mutations being causal to the *tornado* phenotype can be effectively ruled out given the number of phenotypically identical independent *tet1* mutants (Cnops et al., 2006). All previously described *tet1* alleles harbour mutations that disrupt the *TET1* open reading frame in or after the large extracellular loop between transmembrane domains 3 and 4 (Cnops et al., 2006). Therefore, the existing *tet1/trn2* alleles might produce partial *TET1* transcripts, and the resulting truncated TET1 proteins could cause the *tornado* phenotype in weakly semidominant-negative manner, which could have been easily mis-interpreted as recessive loss-of-function. To test this hypothesis, we cloned fragments of *TET1* mimicking the *trn2-1* and *trn2-7* alleles and drove them from the strong constitutive *35S* promoter in Col-0 background. Both p*35S::TRN2-1* and p*35S::TRN2-7* constructs indeed yielded plants with the *tornado* phenotype (**Figure 2**), albeit only partially penetrant. Surprisingly though, we observed the *tornado* phenotype also among plants harbouring the full-length p*35S::TET1g* construct, as well as p*35S::TET1-GFP* and p*TET1_3600_:TET1g-TurboID-HA* (**Figure 2** and **Figure S2**). We further confirmed that the p*35S::TET1g* construct indeed resulted in *TET1* overexpression by qRT-PCR and, despite that we also observed lower levels of endogenous *TET1* expression, there was no correlation between the amount of endogenous or transgenic *TET1* transcripts and the penetrance of the *tornado* phenotype in a population of independent p*35S::TET1g* T1 transformants (**Figure S2**). This result argued against knock-down of endogenous *TET1* due to co-suppression (Depicker and Van Montagu, 1997) causing the tornado phenotype in p*35S::TET1g* plants. Collectively, these results suggested that ectopic expression of full-length, truncated or tagged versions of *TET1* cause the *tornado* phenotype.

**Figure 2:**
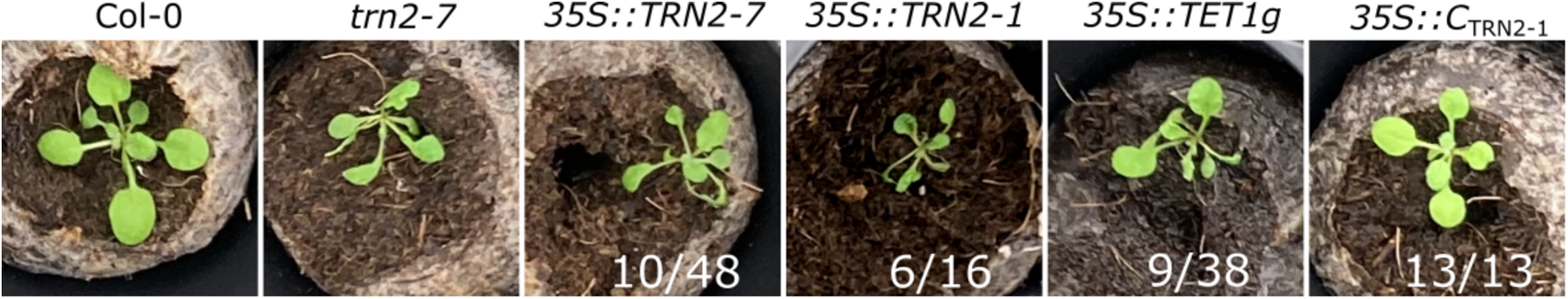
Phenotype of *TET1* full-length and truncated gain-of-function lines. Representative images showing phenotypes of 7 week old plants, specifically the rosette. Left to right: Col-0, *trn2-7*, and Col-0 transformed with the indicated constructs. Ratio’s represent the number of plants with the corresponding phenotype.

To decisively uncouple the effects of loss of *TET1* function from potential dominant-negative effects of residual truncated TET1 proteins, we utilized the CRISPR-Cas9 technology to produce a *bona fide tet1* full knockout or deletion mutant. By co-expressing Cas9 with gRNAs targeting the *TET1* 5’ and 3’ UTRs, we were indeed able to isolate a mutant with the *TET1* coding region cleanly excised and named it *tet1-cko1* (Crispr KnockOut 1) (**Figure 3A-E**). The overall seedling and rosette stage phenotype of the *tet1-cko1* mutant was indistinguishable from the previously described *tet1/trn2* alleles (**Figure 3B,D** and **Figure S3**). We next analysed vascular cell file numbers of *tet1-cko1* roots and observed a similar reduction compared to wildtype as in the *trn2-1* and *trn2-7* alleles (**Figure 3F,G**). Among other defects, previously described *tet1/trn2* mutants often lack one or more epidermal cell files which are replaced by ectopic lateral root cap cells, presumably due to defects in cell division orientation of the protoderm initials (Cnops et al., 2000; Fendrych et al., 2014). Notably, in ∼20% *tet1-cko1* roots, we observed analogical defects in the ground tissue, specifically missing endodermal cell files, resulting in cortex cells directly neighbouring with pericycle cells (**Figure 3F**).and Collectively, these results strongly suggest that the *tornado* phenotype of the previously described *tet1/trn2* alleles is indeed caused by loss of *TET1*. Nonetheless, they also show that gain-of-function of full-length, truncated or tagged versions TET1 can phenocopy *tet1* loss-of-function in a dominant-negative manner, which could explain why we were not able to complement the *tet1* alleles *trn2-7* and *trn2-1*. We thus concluded that the *tet1* deletion allele *tet1-cko1*, which phenocopies other *tet1/trn2* alleles and displays an additional defect in ground tissue patterning and is thus a superior genetic background for further functional characterization of TET1.

**Figure 3:**
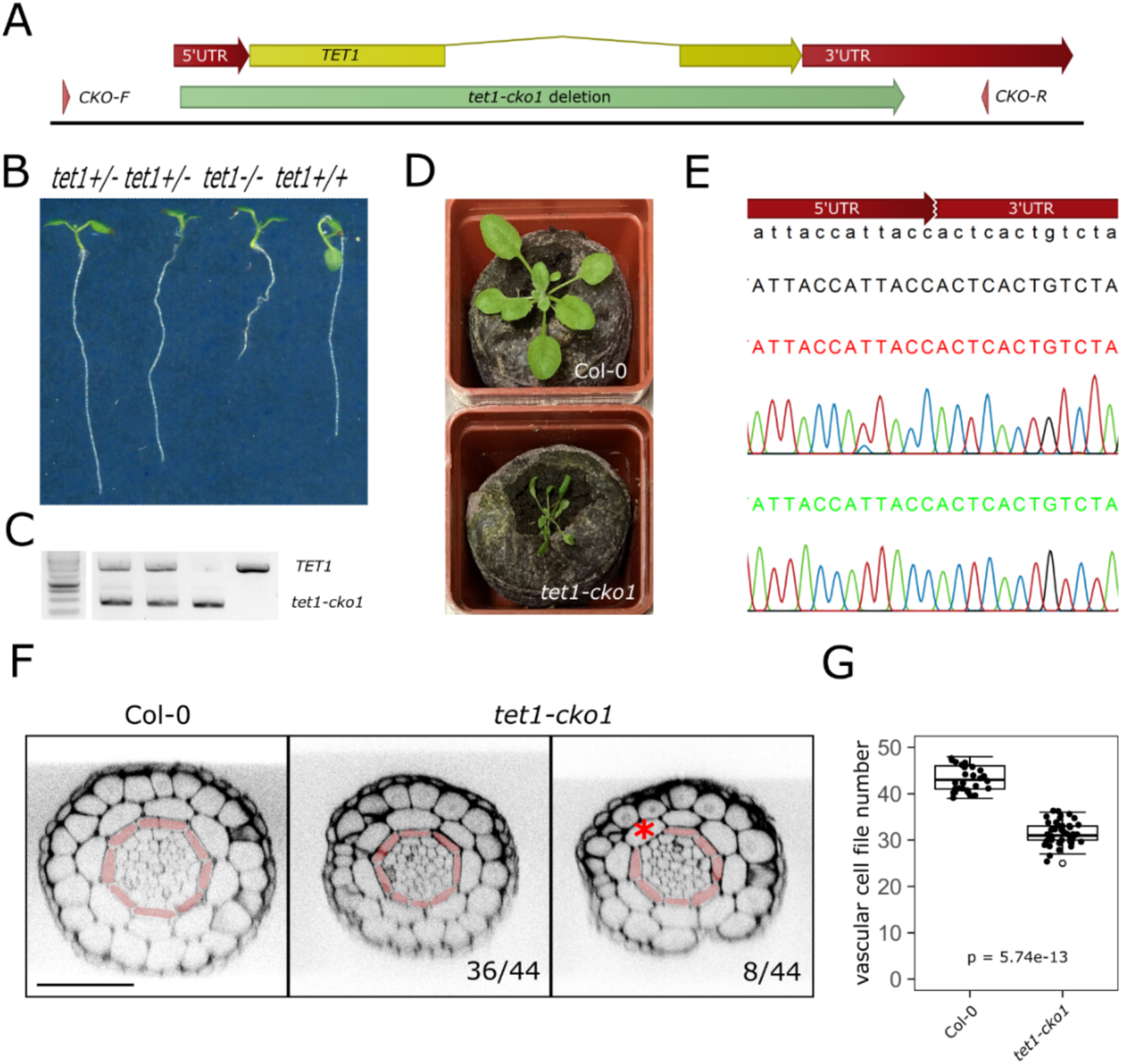
Generation and characterization of *tet1-cko1*, a full knock-out allele of *tet1*. **A**) Schematic overview of the *TET1* coding region and area of deletion by CRISPR. Red represents the *TET1* UTRs, yellow the *TET1* genomic coding region, green the area of deletion by CRISPR and the arrowheads the primer binding region. **B**) Representative images of 7 day old seedlings grown on ½ MS-media. Genotypes, left to right: *tet1*^+/-^, *tet1*^+/-^, *tet1*^-/-^, *tet1*^+/+^. **C**) Genotyping seedlings from panel b. For these DNA bands, *CKO-F* and *CKO-R* primers were used, shown in panel A. Benchtop 1kb ladder shown on left. **D**) 7 week old Col-0 and *tet1-cKO* plants. **E**) Chromatogram showing the *tet1-cKO* DNA sequence spanning the deleted *TET1* coding site. **F**) Representative images of transverse cross-sections of PS-PI stained meristematic roots of Col-0 and *tet1-cKO,* used for quantification shown in panel G. Red colouring represents endodermis. 36/44 roots show a decreased amount of cells in the vascular bundle, while 8/44 additionally show a missing endodermal cell file (indicated with an asterisk). Scale bar = 50µM. **G**) Quantification of the PS-PI treated Col-0 and *tet1-cKO* roots. n=30

### Subcellular localization supports a function for TET1 in ground tissue patterning

We next re-assessed the functionality of TET1 fluorescent fusions by complementing the *tet1-cko1* deletion mutant. As we have had issues complementing the *trn2-7* allele with tagged constructs, we initially decided to use the non-tagged *pTET1*::TET1g construct to look for complementation in the *tet1-cko1* mutant background. We observed full rescue of the *tet1-cko1* mutant by the non-tagged *pTET1*::TET1g construct, but similarly to the *trn2-7* allele, no rescue by p*TET1:*:TET1g-mCitrine, p*TET1*::TET1g-mScarlet or p*TET1*::mCitrine-TET1g (**Figure S3A-D**). As these constructs consisted of identical promoter and coding regions and differed only in the presence/absence of the fluorescent tags, we conclude that the lack of rescue of was indeed caused by non-functionality of the fusion proteins, rather than discrepancies between the endogenous and transgenic expression pattern and/or levels. Difficulties in obtaining functional translational reporters of transmembrane proteins caused by dysfunctionality of N-or C-terminal fluorescent fusions have been reported before (Abas et al., 2006; Wisniewska et al., 2006; Xu and Scheres, 2005). In case of the PIN-formed (PIN) auxin efflux carriers, this obstacle was successfully circumvented by inserting the fluorescent protein into the central hydrophilic loop, or by terminal fusion to the hemagglutinin epitope (HA) tag, which is much smaller than the relatively large fluorescent proteins (Abas et al., 2006; Ntoukakis et al., 2011; Wisniewska et al., 2006; Xu and Scheres, 2005). Analogically, we inserted the mCitrine tag into the single short intracellular loop within the predicted TET1 structure. Nonetheless, this in-loop fusion protein did not complement the *trn2-7* mutant, displayed cytoplasmic localization in T1 generation and got silenced by T2, and caused the *tornado* phenotype in a dominant-negative manner similarly to the p*35S*::TRN2-7 and p*35S*::TRN2-1 constructs, indicating it was not functional either (**Figure S4**). Next, we tagged TET1 N-and C-terminally with three repeats of the HA tag. The p*TET1*:: 3xHA-TET1g (HA-TET1) construct yielded *tet1-cko1^-/-^* T1 plants with wildtype-like phenotype which produced viable T2 seeds, and thus fully rescued the *tet1* loss-of-function mutant, as further confirmed in T2 and T3 generations (**Figure S3E**). These results indicated that the HA-TET1 fusion protein is functional and finally allowed us to reliably assess the subcellular localization of TET1. The expression pattern of p*TET1*:: 3xHA-TET1g was consistent with the scRNA-seq data and the p*TET1*::TET1g-sYFP reporters, once again confirming that the lack of rescue by the fluorescent reporters was due to non-functionality of the fusion proteins rather than incorrect expression of the transgenes (**Figure 4A**). On the subcellular level, the functional HA-TET1 fusion protein displayed dual localization at the PM and in the endomembrane system, the PM signal being more pronounced in younger cells and in the xylem (**Figure 4A,B**). Upon treatment with the ARF-GEF inhibitor Brefeldin-A (BFA), the HA-TET1-positive endomembrane compartments aggregated into the so-called BFA bodies together with PIN1 (Lam et al., 2009) (**Figure S5A**). These results indicate that endomembrane trafficking of HA-TET1 occurs along BFA-sensitive secretory and/or recycling pathways. The PM pool of HA-TET1 was distributed uniformly across the individual PM domains of the meristematic and transition zone cells (**Figure 4B**). This contrasted with the subcellular localization of TET1-sYFP, which was often enriched at the centres of individual apical/basal PM domains of cortex and epidermis cells when expressed from the endogenous p*TET1* or strong meristematic p*RPS5A* promoters (**Figure S5B**), reminiscent of the subcellular localization of the receptor-like kinase IRK in Cortex-Endodermis Initials (CEIs) (Campos et al., 2019) the choline transporter CHER1 in the phloem sieve tubes (Dettmer et al., 2014), or the “super-polar” distribution of PINs (Kleine-Vehn et al., 2011)(**Figure S5**). Nonetheless, as HA-TET1, but not TET1-sYFP, rescued the *tet1* mutant, we favour the hypothesis that the “super-polar” distribution of TET1-sYFP was an artifact caused by incorrect folding, ectopic interactions of the fusion protein and/or simply polar secretion visualized by the dysfunctional fusion. The uniform PM distribution of HA-TET1 was a better proxy of the endogenous TET1 localization. Notably, in the CEIs, CEI daughters (CEIDs) and young cortex and endodermis cells, HA-TET1 localized almost exclusively at the PM and its distribution was often asymmetric, with signal maxima at the apical, basal and/or lateral PM domains (**Figure 4C**). Together with the missing endodermis cell file phenotype of the *tet1-cko1* mutant (**Figure 3F**), this result suggests that TET1 is involved in the control of ground tissue patterning, presumably by linking cell polarity cues to division plane positioning in CEIs and/or CEIDs.

**Figure 4:**
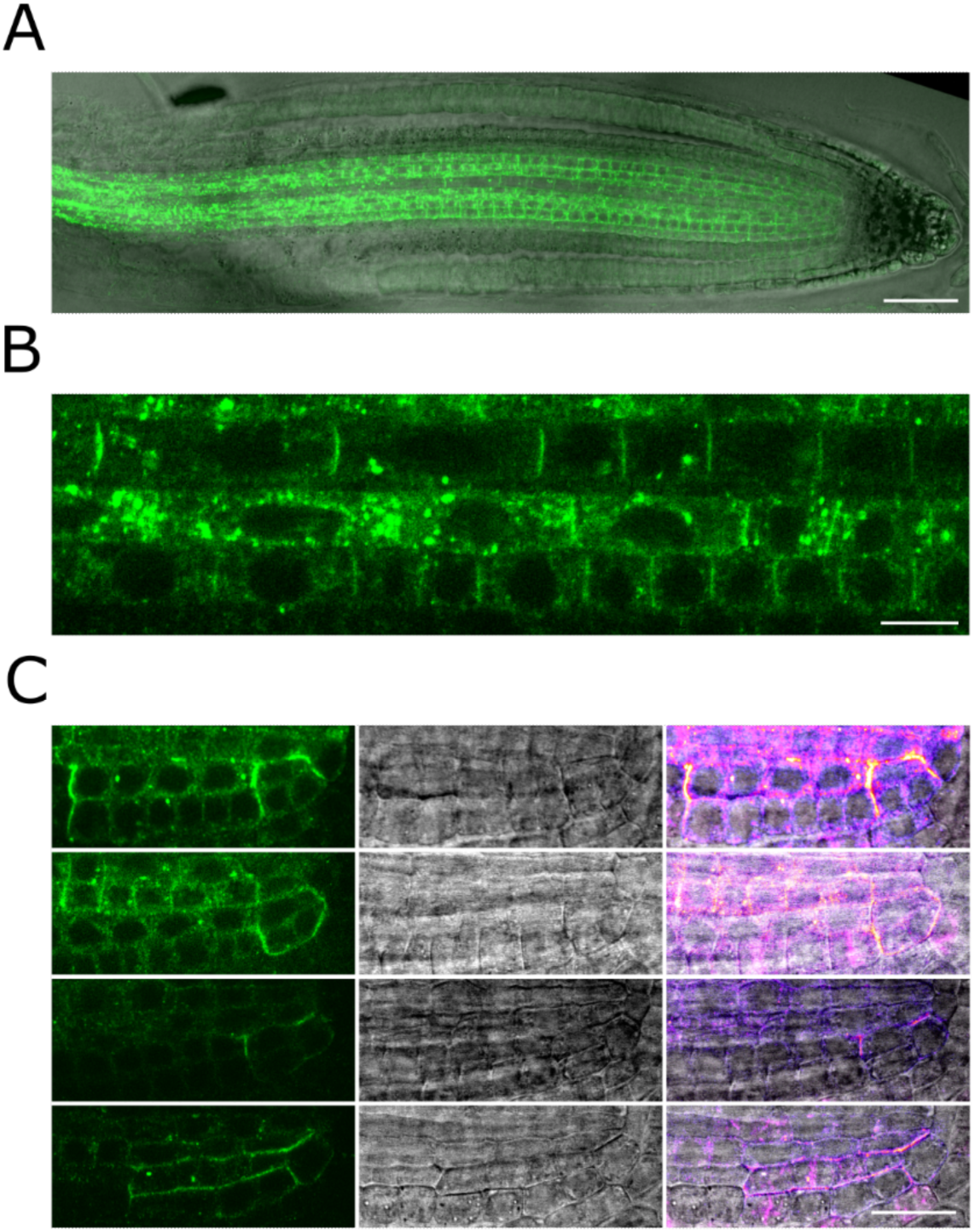
Subcellular localization of HA-TET1. **A**) Confocal image of *pTET1:: 3xHA*-*TET1g* localization *in the tet1-cko1* background, with bright field overlay. Scale bar = 50 μm. **B**) HA-TET1 localization in vasculature, with pronounced membrane localization in the xylem file and more intracellular signal in surrounding procambium cells. Scale bar = 20 μm. **C**) HA-TET1 localization in the CEI, CEIDs and young cortex and endodermal cells. Scale bar = 20 μm.

Taken together, the *TET1* expression pattern; the mutant phenotype of the newly generated *tet1-cko1* deletion allele; and the subcellular localization of the functional HA-TET1 fusion protein all support a role for TET1 in the regulation of cell division orientation during vascular proliferation and ground tissue patterning.

## Discussion

A major unanswered question at the interface of plant cell and developmental biology is how tissue and cell polarity cues are integrated into precisely positioning the cell division plane (Glanc, 2022). In order to find novel regulators of this process in the context of developing vascular cells of the Arabidopsis root, we took a transcriptomics approach to pinpoint those factors being expressed at the correct spatiotemporal location. Based on overlap analysis of transcripts enriched in young phloem and procambium with those upregulated upon ectopic activation of the TMO5/LHW-DOF2.1 pathway (De Rybel et al., 2013; Smet et al., 2019), we identified TET1/TRN2, a member of the evolutionary conserved tetraspanin protein family (Reimann et al., 2017) as a promising candidate of vascular cell division orientation. By analysing the promoter region of *TET1*, we found that the expression domain depends more on regulatory elements in the intron compared to the length of the promoter and, most importantly for its function, that *TET1* is expressed in vascular cells as was predicted by single-cell transcriptomics (Wendrich et al., 2020). Moreover, *trn2-7* and *trn2-1* mutants showed a significant reduction in the number of vascular cell files in the root meristem, supporting a role of TET1 in the controlled switch from anticlinal to periclinal/ radial division plane positioning during vascular development. To prevent potential side effects of truncated TET1 proteins in existing mutants, we generated a full knock-out TET1 mutant, *tet1-cko1*. Besides reiterating the reduction of vascular cell file numbers in the root meristem as the previously described alleles, an additional novel phenotype in the ground tissue was found, with a proportion of the mutants missing an endodermal cell file. These results further support the notion that TET1 is crucial for cell division orientation and reveals an exciting new link to ground tissue patterning. To further characterize TET1, we generated and analysed protein localization in the *tet1-cko1* mutant complemented with the functional HA-TET1 construct. This revealed TET1 localization at the plasma membrane and endomembrane system of the root vasculature and asymmetrical polar distribution at the CEI and young cortex and endodermal cells. Our study has thus established TET1 as a regulator of vascular proliferation and also revealed the involvement of TET1 in ground tissue patterning, providing the necessary foundation for future work of understanding the molecular mechanisms behind these phenomena.

Besides these biological insights, our work again demonstrates the importance of mutant complementation and careful analysis of generated plant material. Cross-checking expression profiles with publicly available single cell datasets allows validation of reporter line expression profiles, as is the case with *TET1,* where meristem expression depends on the presence of the intron. Initially, we aimed to complement *trn2-7* and *trn2-1* mutants with fluorescent protein fusions. After screening a multitude of constructs, we did not find a single one that could rescue the *tet1/trn2* phenotype (Cnops et al 2006). As all 7 of the previously characterized TET1 mutant alleles have mutations around the large extracellular loop (Cnops et al 2006), we investigated whether our failure to complement these mutants is because of a dominant-negative effect due to possible expression of truncated protein. We were surprised to find the *tet1* phenotype present not only in Col-0 lines carrying truncated versions of TET1, but also in overexpression lines harbouring full length TET1 and ruling out potential silencing issues. These results demonstrate that the *tet1* phenotype can be caused by both gain-of-function and loss-of-function and argues that a precise balance of TET1 activity is necessary for plant development. We finally managed to complement a true full-deletion mutant, *tet1-cko1,* with the *pTET1::TET1g* and *pTET1::HA-TET1g* constructs, but failed to do so by *pTET1::TET1g-mCitrine*, *pTET1::mCitrine-TET1g* or *pTET1::TET1g-HA*, which differed only in the type and position of the tags. This result confirmed that fluorescent C-terminal, as well as N-terminal, TET1 fusions are not functional. This necessitates careful re-evaluation of any data involving such reporters, as well as well as fluorescent fusions of other TET homologues whose functionality had not been tested (Yamaguchi et al., 2017; Zhu et al., 2022). Overall, we hope that our work demonstrates the importance of generating full knock out mutant lines and checking for full complementation before continuing any sort of in-depth analysis of generated material, as well as providing a solid foundation for further work on TET proteins in root development.

## Acknowledgements

The authors thank Veronique Storme for assistance in data analysis, Thomas Eekhout for help with data submission, Mieke van Lijsebettens for sharing unpublished constructs and lines and critical reading of the manuscript, and the VIB Bioimaging core and the IEB ASCR Imaging facility for imaging assistance. M.G. is grateful to Matyáš Fendrych for hosting him in his laboratory during the final stages of the project, and to all members of the Fendrych lab as well as Roman Pleskot and Nemanja Vukasinovic for valuable discussions.

## Author Contributions

M.G. and B.D.R. conceived the project; B.D.R., M.G., E.M. and N.K. designed experiments and analysed data; E.V., J.N., K.V., E.M., N.K. and M.G. performed experiments; D.V.D., F.W. and M.V.L. contributed to data interpretation and revising the manuscript; B.D.R. and M.G. supervised the project; M.G., N.K. and B.D.R. wrote the paper with input of all authors.

## Data availability

All quantitative data supporting the findings of this study and respective statistics are available as Table S5. Materials and resources of this study are available from the corresponding author upon reasonable request. Raw data for the bulk RNA-seq analysis can be accessed at NCBI with GEO number: GSE244902.

## Materials & Methods

### Bulk RNA sequencing and data analysis

The *proRPS5A::DOF2.1-GR* construct was cloned using the Multisite Gateway cloning system. 5-day-old seedlings were grown on half strength MS medium plates and then transferred to plates supplemented with 10µM Dexamethasone (DEX). The DEX treatment was performed for 1-2-4 hours before harvesting the bottom 0.5cm of ∼300 roots and three biological repeats per time points were used. Roots were harvested directly into liquid nitrogen and RNA was extracted with the RNeasy kit (QIAGEN). The *Arabidopsis thaliana* Col-0 TAIR10 Genome data was used for mapping and annotation. The analysis was performed with R software package edgeR (R version 3.5.1).

Genes with expression values higher than 0.35 cpm (5 read counts) on at least 3 samples were retained for the analysis. TMM normalization was applied using the calcNormFactors function. Variability in the dataset was assessed with a MDSplot. The treatment and genotype factors were collapsed into one factor. Trended negative binomial dispersion parameters were estimated based on an additive no intercept model with the collapsed factor and a batch effect using the estimateDisp function and down-weighting outlying genes. A quasi likelihood negative binomial regression model was then used to model the over dispersed counts for each gene separately as implemented in the function glmQLFit. The interest was in the simple tests of effect: testing for a treatment effect in the genotype and testing for a genotype effect under treatment conditions. Contrasts were estimated using empirical Bayes quasi-likelihood F-tests. P-values were corrected using FDR method described by Benjamini and Hochberg (1995) for each contrast separately. All edgeR functions were applied with default values. A gene was called differentially expressed when the FDR value was < 0.05.

### Data overlap analysis

From the *proRPS5A::DOF2.1-GR* dataset, genes with fold change >1.5 across all timepoints and p-values < 0.001 across all timepoints were retained. From the published scRNA-seq dataset (Wendrich et al., 2020), procambium and early phloem cell clusters were selected and those differentially expressed genes showing high expression in these clusters (pct.1>0.99) were retained. Overlap analysis of these gene lists was performed using an on-line Venn diagram tool (https://bioinformatics.psb.ugent.be/webtools/Venn/).

### Plant material and growth conditions

*Arabidopsis thaliana* Columbia-0 (Col-0) background was used for generation of mutant and transgenic lines. Seeds were sterilized using 75% EthOH and 25% bleach for 5 minutes, after which they were rinsed twice with 70% EtOH and once with 96% EtOH. Seedlings were grown at 22°C under continuous light on ½ Murashige & Skoog (MS) with 1% agar without sucrose, after a stratification period of 48 hours. 7 DAG seedlings were placed on Jiffy peat pods for phenotypic analysis at rosette stage and propagation. The *trn2-1* (Cnops et al., 2000) and *trn2-7* (GK-254G01.02) (Wang et al., 2015) mutants were described previously. The *tet1-cko1* mutant and all transgenic lines described in this study were generated by transforming the respective constructs into *Col-0*, *trn2-7^+/-^* or *tet1-cko1^+/-^*using the floral dip method (Clough and Bent, 1998). The AGI identifiers for the genes used in this study were as follows: *DOF2.1*: *AT2G28510*; *TET1*: *AT5G46700; UBC: AT5G25760; EEF1α4: AT5G60390*.

### Molecular cloning

All primers and vectors used in this study can be found in **Table S3** and **Table S4**, respectively. Promoters and coding sequences were commercially synthesized (Eurofins) or PCR-amplified using the Q5 high fidelity polymerase (NEB) according to manufacturer’s instructions. Constructs were generated by Multisite Gateway (Karimi et al., 2007) or the Golden Gateway (Karimi and Jacobs, 2021) methods. All entry clones were verified by Sanger sequencing. All sequence data were handled using CLC Main Workbench Version 21.0.4.

### CRISPR/CAS9 mutant generation

To generate full knock out *TET1*, the CRISPR-CAS9 technique was used as described previously (Decaestecker et al., 2019). Briefly, gRNA sequences targeting the *TET1* 5’ and 3’ UTRs were designed using the Cas-Designer tool (Park et al., 2015) and cloned into the *pFASTRK-AtCas9-AG* destination vector. The *TET1-cKO* construct was transformed into Col-0 and candidate genome-edited T1/M1 plants were identified by PCR using primers flanking the gRNA target sites. The excision of *TET1* was confirmed in Cas9-free T2/M2 plants by PCR and Sanger sequencing (**Figure 3**).

### DNA extraction and genotyping

Genomic DNA used for PCR-based genotyping was isolated using the CTAB extraction buffer (0.1 M Tris pH7.5, 0.7M NaCl, 0.01 M EDTA and 0.03 M CTAB) as described previously (Wybouw et al., 2023). Primers used for genotyping can be found in **Table S3**.

### Microscopy and image analysis

GUS staining and imaging was performed as described previously (Wybouw et al., 2023). Fluorescent reporter lines were imaged on Leica SP8 (Leica Microsystems), Zeiss LSM 710 or LSM880 (Zeiss) confocal microscopes equipped with 40x NA 1.1 W (Leica), 20x NA 0.8 or 40x NA 1.2 W (Zeiss) objectives. sYFP and mCitrine were excited at 514 nm and detected at 520-580 nm. mPS-PI staining was done as described previously (Arents et al., 2022). Briefly, 7 DAG seedlings were fixed in 50% methanol/10% acetic acid at 4 degrees overnight, incubated in 1% periodic acid and stained with freshly prepared propidium iodide in Schiff reagent solution for 2 hours or until visibly stained. Seedlings were then mounted in chloral hydrate and imaged in XZ mode on Leica SP8 with a 40x NA 1.1 water immersion objective, excited at 560 nm and detected at 600-700 nm. Immunofluorescence was performed as described previously (Sauer et al., 2006) using the InSitu PRO VSI robot (Intavis) and the following antibodies: mouse αHA (Santa Cruz Biotech) 1:500, rabbit αPIN1 (Baster et al., 2013) 1:1000, AlexaFluor Plus 488-conjugated goat αMouse (Invitrogen) 1:500, AlexaFluor Plus 647-conjugated goat αRabbit (Invitrogen) 1:500. BFA (Sigma) treatment was applied in liquid ½MS medium as indicated in the figure legend from 1000x stock solutions in DMSO. Images were processed and analyzed using FIJI (Schindelin et al., 2012), figures were assembled in Inkscape version 1.1.2.

### RNA isolation and qRT-PCR

Samples from 8 week old Arabidopsis rosettes were snap frozen and ground in liquid nitrogen. RNA was extracted using Trizol and the ReliaPrep RNA Miniprep System (Promega, Netherlands). Trizol was added to each sample for 5 minutes, followed by chloroform. After centrifugation, the top aqueous phase was mixed with isopropanol, incubated for 10 minutes at -20 degrees and then this solution was processed further using the Reliaprep Kit as per the manufacturer’s instructions. cDNA was synthesized from 1 µg of total RNA with the qScriptTM cDNA Supermix kit (Quanta BioSciences, Belgium). qPCR was performed on a Roche Lightcycler 480 with SYBR Green I (Roche, Belgium). Analysis was done with qbase+ Version: 3.3 (https://cellcarta.com/genomic-data-analysis/). Primers used are listed in **Table S3**.

### Quantification and Statistical Analysis

Quantification, statistical analysis and plotting were performed using Microsoft Excel or R version 4.1.0 using RStudio version 1.4.1106 and the ggplot2 package. All data and test used for statistical analyses are summarised in **Table S5**.

